# Somatosensory evoked potentials, indexing lateral inhibition, are modulated according to the mode of perceptual processing: comparing or combining multi-digit tactile motion

**DOI:** 10.1101/2020.10.15.338111

**Authors:** Irena Arslanova, Keying Wang, Hiroaki Gomi, Patrick Haggard

## Abstract

Many perceptual studies focus on the brain’s capacity to discriminate between stimuli. However, our normal experience of the world also involves integrating multiple stimuli into a single perceptual event. Neural circuit mechanisms such as lateral inhibition are believed to enhance local differences between sensory inputs from nearby regions of the receptor surface. However, this mechanism would seem dysfunctional when sensory inputs need to be combined rather than contrasted. Here, we investigated whether the brain can *strategically* regulate the strength of suppressive interactions that underlie lateral inhibition between finger representations in human somatosensory processing. To do this, we compared sensory processing between conditions that required either comparing or combining information. We delivered two simultaneous tactile motion trajectories to index and middle fingertips of the right hand. Participants had to either compare the directions of the two stimuli, or to combine them to form their average direction. To reveal preparatory tuning of somatosensory cortex, we used an established event-related potential design to measure the interaction between cortical representations evoked by digital nerve shocks immediately before each tactile stimulus. Consistent with previous studies, we found a clear suppressive interaction between cortical activations when participants were instructed to compare the tactile motion directions. Importantly, this suppressive interaction was significantly reduced when participants had to combine the same stimuli. These findings suggest that the brain can strategically switch between a comparative and a combinative mode of somatosensory processing, according to the perceptual goal, by preparatorily adjusting the strength of a process akin to lateral inhibition.

## Introduction

Given the overwhelming flux of information and brain’s limited processing capacity (Broadbent, 1958; Luck and Vogel, 1997; Gallace *et al*., 2006), incoming sensory inputs need to be processed efficiently to guide behaviour. Acuity and discrimination thresholds describe the minimal units of sensory information required to identify a sensory input, and have generally been the starting point for characterising sensory systems, both in vision (Watson and Robson, 1981; Westheimer and Wehrhahn, 1994; Schwartz *et al*., 2007) and in touch (Sherrick, 1964; Evans and Craig, 1991; Driver and Grossenbacher, 1996; Soto-Faraco *et al*., 2004; Tamè *et al*., 2011; Rahman and Yau, 2019; Halfen *et al*., 2020). However, our normal perceptual experience of the world is not limited to minimal inputs and minimal contrasts; the brain will often integrate multiple inputs into a single perceptual event. Imagine a flock of birds with each bird moving in a slightly different direction. While an observer can isolate one particular bird’s movement, the observer is also able to perceive the *average* movement of the flock as a whole.

The ability to extract overall or average motion information from multiple, simultaneous motion cues has been described in vision (Watamaniuk *et al*., 1989; Watamaniuk and McKee, 1998) under the idea of *ensemble perception* (for review see Alvarez, 2011; Whitney and Yamanashi Leib, 2018). In touch, a few studies have investigated aggregation of tactile features such as intensity or frequency (Walsh *et al*., 2016; Kuroki *et al*., 2016; Cataldo *et al*., 2019). However, when an object held between fingers begins to move, the overall motion direction of the object can be also clearly perceived (Martin, 1992). Importantly, because the motion cues at each fingertip may not be redundant, the brain must aggregate individual motion direction cues from different digits to extract the veridical average motion direction. The cognitive and physiological mechanisms of such ensemble perception for spatial aspects of touch remain unclear.

In contrast, more effort has been made to understand the mechanisms that support acuity and discrimination. In particular, lateral inhibition – a pervasive neuroanatomical principle of sensory system organisation – has been found to sharpen discrimination via local networks of inhibitory interneurons. Briefly, inhibitory interneurons connect adjacent cortical neurons so that firing of one cortical neuron tends to lead to inhibition of its neighbours. This arrangement enhances responses to small spatially-detailed stimuli relative to spatially-extended stimuli, since the former do not trigger lateral inhibition from neighbouring receptive fields (RFs), whereas the latter do. This general principle has been confirmed by neurophysiological studies of neurons in visual (Blakemore and Tobin, 1972; DeAngelis *et a*l., 1992; Angelucci *et al*., 2017), olfactory (Urban, 2002), auditory (Foeller *et al*., 2001; Wehr and Zodor, 2003; Kato *et al*., 2017), and somatosensory (Laskin and Spencer, 1979; Brumberg *et al*., 1996; Dykes *et al*. 1984; DiCarlo *et al*. 1998; Brown *et al*., 2004; Mirabella *et al*., 2001; Sachdev *et al*., 2012) cortices.

However, previous studies have focused on very local interactions within a single digit. In these cases, lateral inhibition is thought to sharpen RF tuning, thus increasing spatial acuity (Brown *et al*., 2004; Haggard *et al*., 2007; Cardini *et al*., 2011) and enhancing contrast (Brumberg *et al*., 1996). Yet, sensory representations of *individual digits* in primary somatosensory cortex (SI) are also laterally connected via inhibitory (but also excitatory) interneurons (Forss *et al*., 1995; Reed *et al*., 2008). Indeed, inter-body lateral connections seem to underlie the very rapid spread to adjacent body parts of the digit RF of SI neurons, when the digit forming their original RF is surgically amputated (Calford and Tweedale, 1991; Kelly *et al*., 1999; Foeller *et al*., 2005). Therefore, the role of these longer-range (inhibitory) lateral connections is shaping RFs and maintaining topographic organisation of cortical maps. Moreover, they have been also shown to modulate tactile judgements that require integration of information across adjacent fingers (e.g., Wilimzig *et al*., 2012).

Lateral inhibitory interactions in vision also spread across distances greater than the RFs of adjacent primary visual neurons (Fitzpatrick, 2000; Mareschal *et a*l., 2010). Such lateral inhibition is believed to contribute to a visual phenomenon called “repulsion”, which manifests as an exaggeration of contrast between two visual stimuli (Solomon, 2020). For example, in a classical tilt illusion (Gibson, 1937; Clifford, 2014), “repulsive bias” can reflect the exaggeration of the difference between the orientations of neighbouring Gabor patches, so that the target Gabor’s tilt is biased away from the orientation of flanker Gabors. Interestingly, Mareschal and colleagues (2010) showed that such repulsion occurs even when the distance between target and flankers exceeds the size of individual RFs, suggesting that lateral inhibition spreads across neighbouring RFs, exerting inhibitory influence on more distant neurons.

Considering that lateral inhibition acts to amplify contrast and that tactile connections can span adjacent digits, lateral inhibition would seem particularly relevant for tasks requiring precise localisation of stimuli within a finely-tuned topographic map, such as detecting differences between stimuli concurrently applied to adjacent fingers. In contrast, such inhibition would seem dysfunctional when concurrent sensory inputs from adjacent digits need to be combined to compute an overall percept. This is because adjacent representations will tend to interfere (Harris *et a*l., 2001; Tamè *et al*., 2014), making it difficult to form precise and accurate representations of each stimulus, and thus distorting the aggregated final percept. For example, blocking lateral inhibition in the fruit fly’s visual system reduces encoding of an individual object’s movement direction, but increases encoding of overall pattern motion (Keleş and Frye, 2017). In a similar vein, excessive lateral inhibition may hinder integrative processing (Bertone *et al*., 2005; Gustafsson, 1997).

In general, perceptual processing sometimes needs to identify what differs between multiple simultaneous stimuli, and at other times needs to identify what multiple stimuli have in common. Strategically regulating the degree inhibition between representations from different regions of the receptor surface might offer one method to achieve this perceptual flexibility. We therefore sought to examine whether the brain can *strategically* regulate the strength of inhibition between multiple stimulus representations. Such a process might potentially allow the brain to implement two distinct perceptual modes when processing inputs from adjacent regions of the receptor surface, either comparing or combining information as appropriate.

Lateral inhibition is often demonstrated by visual phenomena such as Mach bands or by masking effects in touch. However, non-invasive studies of the mechanisms underlying lateral inhibition in human somatosensation are scarce. A number of studies have suggested an inhibitory mechanism in primary somatosensory cortical areas, which perhaps operates analogously to lateral inhibition in vision. Gandevia and colleagues (1983) found that the cortical potentials evoked by simultaneous stimulation of two adjacent digits had lower amplitude than the sum of potentials evoked by stimulating each digit individually. This underadditive aggregation of evoked responses was linked to (lateral) inhibitory processing (Gandevia *et al*., 1983; Hsieh *et al*., 1995; Ishibashi *et al*., 2000). This suppressive interaction follows the somatotopic receptive field organization (i.e., underadditivity is stronger when stimulation is applied to adjacent skin regions such as index and middle finger relative to index and ring finger; Ishibashi *et al*. 2000; Ferrè *et al*., 2016) and thus cannot be explained solely by response saturation and masking (Severens *et al*., 2010). In addition, this suppression has been found in several regions along the somatosensory pathway, with stronger interactions in the cortex than in brainstem or thalamus (Hsieh *et al*., 1995). Furthermore, such suppressive interactions have been found to vary with the functional state of the sensorimotor system (i.e., following the alteration to the boundaries of cortical sensory maps; Haavik-Taylor and Murphy, 2007). Finally, Cardini and colleagues (2011) showed that the degree of suppression is not fixed, but can be modulated by multisensory context. They observed stronger suppression when viewing one’s own body then when viewing a neutral object in the same location. However, the tactile task was not varied in their study, and always involved acuity judgements for stimuli delivered to the index or middle finger. Thus, to our knowledge, the wider question of how tactile task requirements might influence suppressive somatosensory interactions has not previously been considered.

Here, we investigated whether EEG measures of somatosensory suppression (Gandevia *et a*l., 1983; Hsieh *et al*., 1995; Ishibashi *et al*., 2000) can be modulated as a function of the requirements of a perceptual task. As we have seen, functional perception may involve adjusting the “mode” of neural processing to favour either differentiation between stimuli based on specific details, or synthesising multiple inputs to produce an overall percept. To investigate this possibility, we delivered two tactile motion trajectories, whose spatial directions could differ, simultaneously to index and middle fingertips of the right hand. Participants were asked to either compare the directions of the two stimuli, by reporting the magnitude of the directional discrepancy between them, or to combine the two stimuli by reporting their average direction.

Averaging two distinct sensory cues is a form of cue integration. However, while the main aim of cue integration is to form a maximally reliable new percept (Ernst and Bülthoff, 2004), the main aim of averaging is to extract overall gist information (Whitney and Yamanashi Leib, 2018). In an averaging task, optimal performance requires participants to allocate equal weights to both cues. Suppressive interactions between finger representations would be dysfunctional for averaging. In contrast, because lateral inhibitory mechanism has been suggested to amplify difference (Solomon, 2020), it would be beneficial for detecting small differences between stimuli applied to adjacent fingers.

The interaction between cortical representations of the stimulated digits was measured immediately before presentation of tactile stimuli that participants either compared or combined. This allowed us to probe the preparatory tuning of inhibitory mechanisms. We predicted that the task instruction would lead to strategic top-down modulation of the state of suppressive interactions, in expectation of either comparing stimuli, or combining them. In other words, the brain might prepare the appropriate mode of processing in advance of stimulation, by tuning circuits in somatosensory cortex accordingly.

## Methods

### Participants

15 right-handed volunteers (aged 20 to 39 years with mean age of 24.5, 9 women), who were naïve to the paradigm and research questions, took part in the experiment. Because no previous study, to our knowledge, has reported an effect size estimate for the effect of tactile task on EEG measures of somatosensory suppression, we could not perform a formal power calculation. However, one previous study has reported multisensory modulation of somatosensory suppression (Cardini *et al*., 2011), obtaining large effect sizes with *n* = 15. While one might assume that task-related modulations of somatosensory suppression would be similar in strength to multisensory modulations, we could not find firm grounds for that assumption in the existing literature. In the absence of more specific information, we therefore followed the sample size used by Cardini et al., (2011). All participants reported normal or corrected-to-normal vision and no abnormalities of touch. They all provided a written consent. Procedures were approved by the University College London (UCL) research ethics committee and were in accordance with the principles of the Declaration of Helsinki.

### Tactile apparatus

The tactile apparatus consisted of two spherical probes (4 mm diameter) attached to two stepper linear actuators (*Haydon Kerk Motion Solutions* 15000 series, model LC1574W-04) that were fixed to two motorised linear stages (Zaber X-LSM100B, Zaber Technologies Inc., Canada) mounted in an XY configuration (Figure 1C). The actuators were controlled by a microcontroller (Arduino) and were moved up and down to let the probe make static contact with the skin at the start of tactile stimulation and retract after the end of stimulation. The linear stages were controlled by custom Matlab scripts that allowed the probe to be moved in predefined trajectories (see *Tactile stimuli* below). The apparatus was covered by a box with a small aperture. The to-be-stimulated right index and middle fingertips were positioned over the aperture, and secured with foam padding. Participants rested their right hand in a fixed palm-down position, so that, through the aperture, the probes lightly touched the centre of their index and middle fingertips. A webcam was placed under the apparatus to monitor the finger placement and contact with the probe. The distance between the fingers was approximately 25 mm. The hand was then covered with a screen, so that the probes could not be seen.

**Figure 1.**
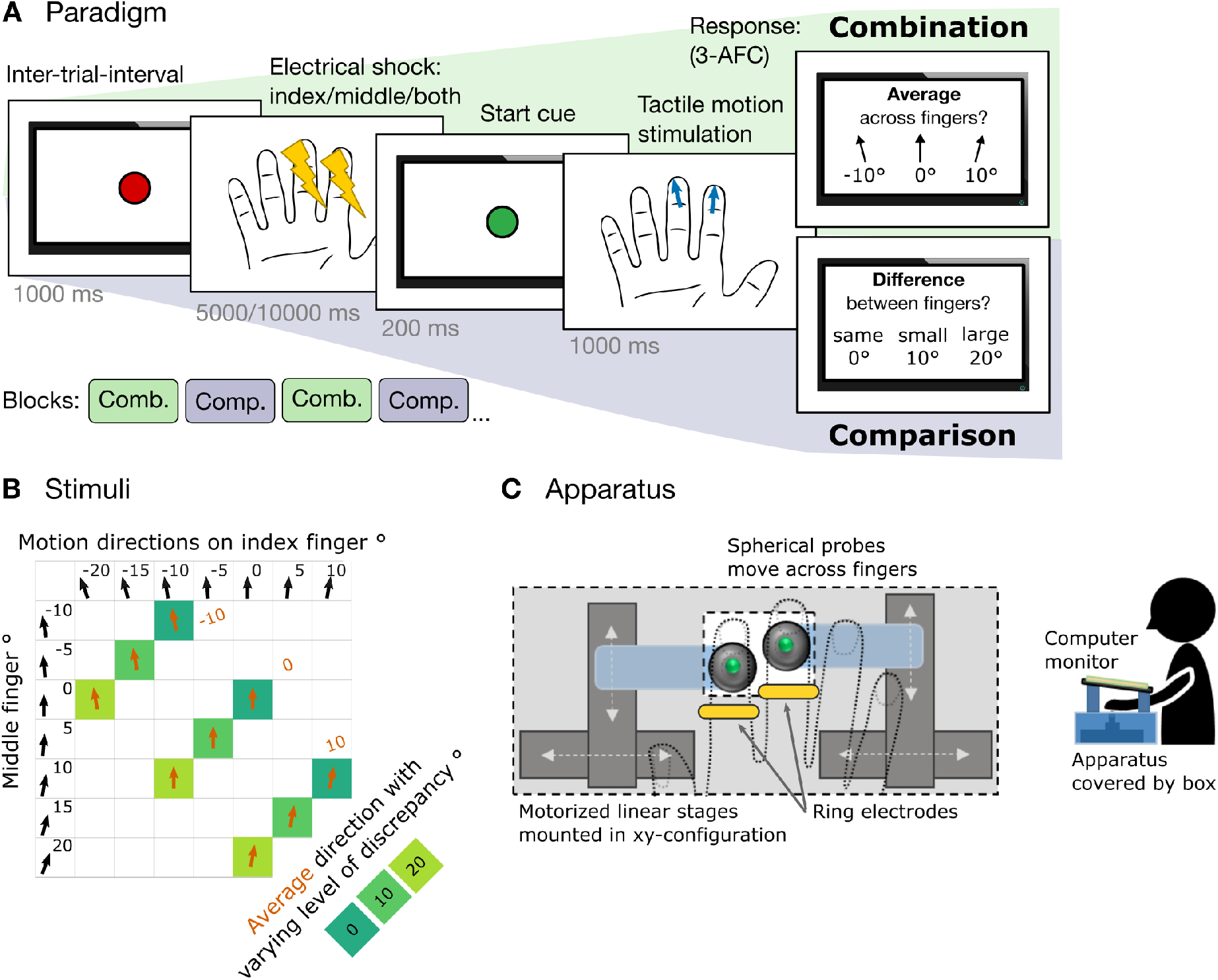
Experimental paradigm and setup. (A) Participants (n = 15) performed two different perceptual tactile motion tasks in alternating blocks. In combination task, participants averaged two tactile motion trajectories, whereas in comparison task, they discriminated between the trajectories. Prior to tactile motion stimuli, mild digital nerve shocks were delivered to the to-be-stimulated fingers to reveal preparatory somatosensory-evoked activity. (B) Nine different pairs of tactile motion stimuli produced three average direction patterns (−10°, 0°, 10°) with three levels of discrepancy (0°, 10°, 20°). Stimuli were identical in both tasks. (C) Motorised linear stages produced continuous tactile motion along the fingertips. The apparatus was covered by a box with a small aperture. To-be-stimulated fingertips were positioned over the aperture, and secured with foam padding. The aperture and the hand were then covered with a computer screen. Digital nerve stimulation was delivered via a pair of ring electrodes.

### Tactile stimuli: continuous motion and double-finger stimulation

Continuous motion along the fingertips was created by moving the probes at preselected angles ranging from −25 to 25 degrees to the distal-proximal finger axis in 5° steps, at a constant speed of 10 mm/s. The movement of each probe was controlled individually allowing for delivery of trajectories with varying discrepancy simultaneously along both fingertips. Figure 1B shows 9 possible combinations compromised of 7 individual directions delivered simultaneously to two fingers. The combinations produced three different average motion patterns (−10°, 0°, or 10° from straight ahead), with varying levels of discrepancy (0°, 10°, or 20°) between the two stimuli. The duration of each trajectory was approximately 1 s and the distance travelled was 10 mm.

At the beginning of each trial, the probe was advanced to make a static contact with the fingertip. The initial position of the probe was jittered across trials (−2.0, 0.0, or 2.0 mm from the centre of the fingertip) to discourage using memory for locations as a proxy for direction. After each trajectory, the probe was immediately retracted and returned to its starting position. The duration of each trajectory was approximately 1 s and the distance travelled was 10 mm. The sound made by the apparatus was masked with white noise continuously played over headphones.

### Perceptual tasks: comparing and combining double-finger tactile stimuli

Participants performed two different tasks that required them to adapt two distinct perceptual processing modes (comparison or combination; see Figure 1A). The tasks were performed in alternating counter-balanced blocks, four blocks per task. At the beginning of each block, participants were notified which task they were going to perform. The tactile stimulation was identical for both tasks and the mode of processing was manipulated with task instructions. In each trial, after the electrical stimulation, both probes were moved simultaneously along both right index and middle fingertips in pre-specified directions. In comparison task, participants were asked to judge the discrepancy in direction between the pairs of tactile motion trajectories delivered to index and middle fingers. They had to identify whether the discrepancy was zero (0°), moderate (10°), or large (20°). In combination task, participants were required to judge the average direction between the same pairs of motion trajectories. They had to report whether the average direction was more to the left from straight ahead (−10°), straight ahead (0°), or more to the right from straight ahead (10°). The response was given after tactile motion stimulation by pressing a corresponding key with their left hand. Responses were unspeeded, and no feedback was given. In total, participants completed 180 trials per task.

### Digital nerve stimulation

To elicit somatosensory-evoked activity electrical stimulation was delivered via a pair of ring electrodes place over the distal phalanxes of the right index and middle fingers with a cathode 1 cm proximal to the anode, at a rate of 2 Hz. Individual sensory detection thresholds for electrical shocks were determined prior to the main experiment with method of limits. Reversals occurred after participants detected the stimulus twice in a row, resulting in stimulus intensity that corresponded to a 70% detection. Stimulation was delivered with a neurophysiological stimulator (Digitimer DS5 stimulator) as a square-wave pulse current, each pulse lasting 0.2 ms. In the main experiment, stimulation was produced at intensity 1.4 times higher than the individual sensory threshold. In each trial either the index finger, the middle finger, or both fingers were randomly stimulated. Brain activity elicited by stimulating index and middle finger in isolation provided a predicted sum of activity (index + middle) under the assumption of no suppression. If suppression occurred, actual activity during double-stimulation (both) would be reduced compared to the sum of individual stimulations. The electrical stimulation occurred before the tactile motion stimuli to reveal the preparatory tuning of somatosensory cortex. The number of electrical pulses was randomly varied (10 or 20) to make the timing of tactile motion onset partly unpredictable, thereby encouraging participants to maintain preparedness to tactile motion task. In total, there were 900 electrical stimuli delivered for each stimulation condition (index, middle, or both) per task.

### Electroencephalographic (EEG) recording and pre-processing

EEG was recorded from 17 scalp electrodes (Fp1, Fp2, AFz, F3, F4, C5, C3, Cz, C4, C6, CP5, CP3, CPz, CP4, CP6, O1, O2) using a BioSemi ActiveTwo system (BioSemi, 2011). Horizontal electro-oculogram (EOG) recordings were made using external bipolar channels positioned on the outer canthi of each eye. Reference electrodes were positioned on the right and left mastoids. EEG signals were recorded at a sampling rate of 2048 Hz. A trigger channel was used to mark the timing of electrical shocks. Data were preprocessed in Matlab with EEGLAB toolbox (Delorme and Makeig, 2004) and ERPLAB toolbox (Lopez-Calderon and Luck, 2014). Data were re-referenced to the average of the mastoid electrodes, subjected to high-pass (0.5 Hz) and low-pass (30 Hz) filtering. Epochs of 250 ms were extracted spanning from 50 ms before each shock to 200 ms after shock onset. For each epoch, signal between −1 and 8 ms relative to electric shock onset was linearly interpolated in order to remove electrical artifact (Cardini *et al.,* 2011; Cardini and Longo, 2016). Epochs were then baseline corrected to the first 50 ms. Trials with eyeblinks (HEOG left and right channels exceeding ± 80 mV) or with voltage exceeding ± 120 mV at any channel between –50 and 200 ms relative to each shock were eliminated. The mean percentage of trials rejected was 24.1% ± 11.5% in combination task and 23.8% ± 10.9% in comparison task. There were no significant differences in the amount of rejected trials between tasks (*p* = .70) nor between stimulation conditions (*p* = .88). Grand average SEPs were computed separately for the two tasks (comparison and combination) and electrical stimulation conditions (index-alone, middle-alone, both).

### Quantification and statistical analysis

We expected the suppressive effect to arise within the P40 component (Biermann *et al*., 1998; Ishibashi *et al*., 2000; Cardini *et al*., 2011), which reflects the afferent volley and first processing wave within somatosensory cortex. Scalp topographies of P40 showed a positive parietal peak and a reversed polarity over frontal channels (Figure 3A). This reversal across the central sulcus is consistent with prior reports of this component (e.g., Cardini *et al*., 2011), and is a marker of SI processing (Allison *et al*., 1989).

Accordingly, we analyzed the mean SEP amplitudes between 20 to 60 ms following digital shock onset. The time-window was chosen after visual inspection of grand-averaged waveform pooled across all stimulation conditions. Previous studies have tended to see a slightly later onset of the P40 component, starting at 40 ms after stimulation (Cardini *et al*., 2011; Cardini and Longo, 2016; Gillmeister and Forster, 2012). In the present study, the component started slightly earlier around 20 ms after stimulation. We ended our time-window at 60 ms, because from there P40 started to overlap with N70. Thus, we chose the 20 to 60 ms time-window that encompassed the whole component around the peak at 45 ms. The mean SEP amplitudes between 20 to 60 ms were acquired per participant (n = 15) for each stimulation condition (index, middle, both), separately for combination and comparison tasks.

Based on index and middle SEP amplitudes, we calculated the predicted sum under the assumption of no suppression (index + middle). If suppression occurred, SEP amplitudes during double-stimulation (both) would be significantly reduced compared to predicted sum of individual stimulations (under-additivity). Shapiro-Wilk test of normality indicated that all measures did not significantly deviate from a normal distribution (all *p* values were .12 < *p* < .96). Thus the amplitudes were fit into a repeated-measures ANOVA with factors task (combination vs. comparison) and stimulation (both vs. index + middle). Significant interaction would indicate differential somatosensory activation between tasks. The interaction would be followed-up by simple effects analysis comparing stimulation condition across tasks.

To compare suppression between tasks, we calculated a “Somatosensory Suppression Index” (SSI), defined as the difference in amplitude between the arithmetic sum of potentials evoked by two individually stimulated fingers and the potentials evoked by simultaneous stimulation of two fingers (Cardini *et al*., 2011). The SSI was calculated with the following equation:

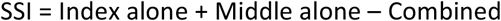

Higher values of SSI indicate stronger suppression within the somatosensory system. A paired-sample t-test was employed to compare SSI between comparison and combination tasks, because there was no significant deviation from normality (Shapiro-Wilk test: *p* = .81).

Behavioural performance was quantified as the accuracy to choose the correct average (in combination task) or correct difference (in comparison task) from three options. Accuracy was then compared across tasks with paired-sample t-test, because there was no significant deviation from normality (Shapiro-Wilk test: *p* = .52).

## Results

### Behavioural performance

After each tactile stimulation, participants were presented with three choices on the computer screen positioned above right hand, and selected one with their left hand (see Figure 1A). Figure 2A shows a confusion matrix with the mean proportion of each response as a function of the actual directional difference (comparison task) or average direction (combination task). Participants performed better in combination task (56% correct, SD = 9%) relative to the comparison task (47% correct, SD = 8%). The difference in performance was significant (paired-sample t-test: *t*_14_ = 3.5, *p* = .004, *d* = .90). Previous studies have also found that somatosensory aggregation tends to produce better performance than discrimination (Cataldo *et al*., 2019), possibly reflecting that the aggregate can be derived even when discrepancy between stimuli is unclear. Performance between the tasks was not correlated across participants (*r* = .20, *p* = .47), showing no evidence for a common computational factor underlying individual differences in performance.

**Figure 2.**
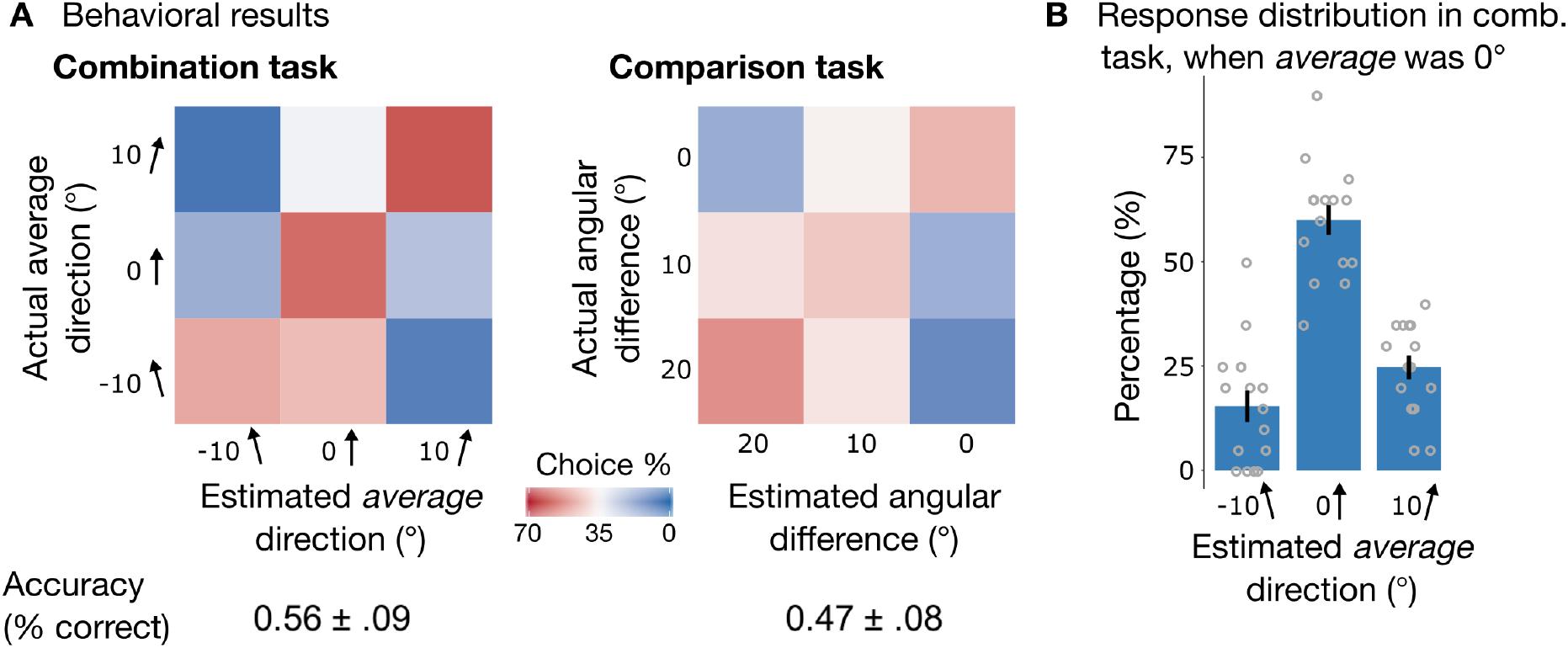
Behavioral results. (A) Confusion matrices illustrate the group-mean percentage of choosing one of three response choices as a function of correct response. Participants were more accurate in combination task (right panel) compared to comparison task (left panel; *p* = .004). (B) Group-mean distribution of responses in combination task, when average direction pattern was straight ahead, but direction on the index finger was to the left (−10°) and direction on the middle finger was to the right (10°).

To ensure that participants truly averaged the discrepant trajectories in the combination task rather than selectively attended to either finger, we plotted the response distribution when true average direction was straight ahead (0°), but direction on the index finger was to the left (−10°) and direction on the middle finger was to the right (10°) (see Figure 2B). Participants correctly identified the average direction to be straight ahead. This indicates that participants tried to combine the two motion directions rather than selectively attended to either finger.

### Somatosensory evoked EEG activity

Figure 3A shows grand mean SEPs (n= 15) elicited by digital shocks immediately before tactile motion stimuli, averaged across electrodes over contralateral somatosensory cortex (C3, C5, CP3, and CP5). Suppression is defined as the amplitude reduction for combined stimulation relative to the sum of the amplitudes for individual finger stimulation. To investigate suppression quantitatively, we first summed the amplitudes for individual index and middle finger stimulations (purple line on Figure 3A). This effectively provides a prediction of the amplitude for combined stimulation under a hypothesis of no somatosensory suppression (i.e., perfect additivity).

**Figure 3.**
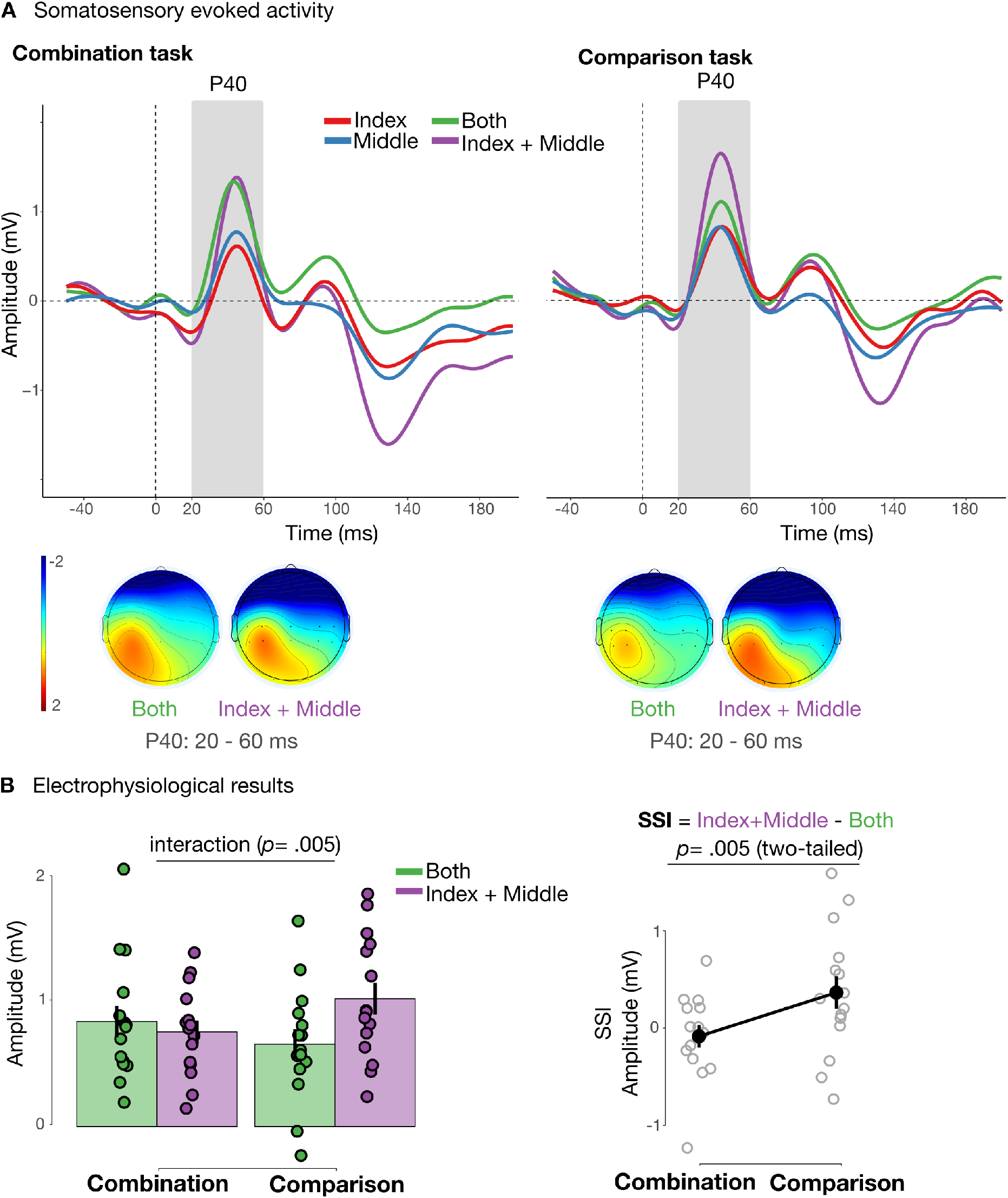
Electrophysiological results: somatosensory evoked potentials and somatosensory suppressive index. (A) ERP waveforms show grand averaged SEPs (n = 15) separately when shocking index finger (red), middle finger (blue), and both (green) fingers simultaneously. In addition, it shows the sum of individual stimulations (purple) that reflects the predicted amplitude for double-stimulation under the assumption of no suppression. The waveforms represent pooled activity across contralateral somatosensory electrodes (C5, C3, CP5, CP3). The grey shaded area shows the analysis time-window that corresponds to P40 component (20 to 60 ms relative to shock onset). Topographic maps show mean activity in the P40 component. (B) Right panel shows mean amplitudes for actual double-shock stimulation (green) and predicted double-shock stimulation under assumption of no suppression (purple) separately for averaging and discrimination tasks. Dots are single participants’ amplitudes and error bars represent SEM. Left panel shows mean calculated SSI (index + middle - both) and its difference between tasks. Grey dots are single participants’ SSI with error bars representing SEM.

We then performed a 2-by-2 repeated measures ANOVA with factors task (combination vs. comparison) and stimulation (both vs. summed-index-and-middle) on the mean amplitudes within a 20 to 60 ms time-window (Figure 3B). The main effects of task (*F*_1, 14_ = .45, *p* = .52, *η*_*p*_^*2*^ = .03) and stimulation (*F*_1, 14_ = 1.23, *p* = .29, *η*_*p*_^*2*^ = .08) were not significant. As predicted, the analysis yielded a significant interaction (*F*_1, 14_ = 10.78, *p* = .005, *η*_*p*_^*2*^ = .44), indicating that the degree of under-additivity varied between the tasks. Importantly, the interaction remained significant after controlling for differences in behavioural performance between the tasks (*F*_1, 13_ = 13.34, *p* = .003, *η*_*p*_^*2*^ = .51), suggesting that the differences in underadditivity between tasks were not simply due to differences in task difficulty. Indeed, given the assumption of a linear relation between performance and SI responses, the true effect of task on somatosensory underadditivity may be *larger* than suggested by the uncorrected means data shown in Figure 3B. We further explored the significant interaction using simple effects analysis. It showed that in combination task, amplitudes to double-stimulation were similar to amplitude to summed-index-and-middle stimulation (*p* = .47). In comparison task, the difference between amplitudes to double-stimulation relative to summed-index-and-middle stimulation became larger (*p* = .47), supporting the predicted shape of the interaction. Simple effects analysis was not Bonferroni-corrected, because it was not used to draw any additional inferences, but merely describe the shape of the significant interaction.

To compare the magnitude of under-additivity between tasks, we calculated the SSI (index + middle – both) separately for comparison and combination task. A 2-tailed paired-sample t-test revealed greater SSI in the comparison task (mean SSI = 0.36 ± 0.65 mV) than in combination task (mean SSI = −0.08 ± 0.44 mV) (*t*_14_ = 3.28, *p* = .005, *d* = .85; Figure 3B). Thus, somatosensory suppressive interactions between stimulated digits were modulated according to the specific perceptual task.

## Discussion

We showed that the suppressive interaction between evoked responses to simultaneous stimulation of two digits was not fixed, but was strategically adjusted according to the perceptual task at hand. When participants compared stimuli on the two fingers, suppressive interaction was stronger than when they combined percepts across both fingers to extract an average. Importantly both comparing and combining require processing information from both digits, so the difference between tasks is not merely in selection or attention. Rather the tasks differed in their post-selection processing. Our results suggest that the neural circuitry of sensory system may be tuned to extract differences in comparison mode, or to extract consistent overall features in combination mode. Switching between these processing modes may involve adjusting the gain of local inhibitory circuits.

In principle, a subadditive interaction could simply reflect a ceiling effect, rather than a specific inhibitory process. For example, the increased stimulus energy in the double shock condition might approach a maximal level of firing in somatosensory neurons. For this reason, our electrical stimuli were kept to low levels. Severens and colleagues (2010) reported a subadditive interaction using a frequency-tagging method, which may avoid some of the interpretational concerns regarding ceiling effects. Finally, even if some saturation similar to a ceiling effect were to be present in our data, we still observed a significant difference between two perceptual tasks in scalp responses evoked by identical stimuli. Thus, ceiling effects alone cannot readily explain our results.

In addition, our results cannot readily be explained by task difficulty. We designed our averaging and discrimination tasks to have comparable levels of performance based on pilot data. However, we did find that discrimination performance in the experiment was significantly worse than averaging performance. In principle, an adaptive design could adjust the stimuli to balance performance across tasks for each participant, to remove this behavioural effect. Even so, the behavioural difference between tasks is unlikely to account for the difference between ERP suppressive interactions, since including performance as a covariate did not abolish (and in fact strengthened) the difference between conditions in suppressive interactions. This argument assumes, of course, a linear relation between performance and somatosensory ERP signal strength. However, this assumption may be reasonable within the supraliminal range studied here.

Studies of tactile perception have historically focused on performance limits for perceiving a single stimulus (Weinstein, 1968; Mancini *et al*., 2014). However, our everyday experience involves dynamic interactions with objects and perception of a single object through multiple skin contacts. Tactile processing bandwidth is too low to perceive multiple independent tactile stimuli simultaneously (Gallace *et al*., 2006). Nevertheless, neurons in some non-primary somatosensory areas (i.e., secondary somatosensory cortex, SII) exhibit large RFs spanning multiple digits, and providing consistent coding of stimulus features, such as orientation, anywhere within the RF (Fitzgerald *et al*., 2006a,b). This implies that the brain can process and integrate information from multiple touches despite limited processing capacity. Yet most previous studies have often focused solely on discrimination between stimuli (Sherrick, 1964; Evans and Craig, 1991; Driver and Grossenbacher, 1996; Soto-Faraco *et al*., 2004; Tamè *et al*., 2011; Rahman and Yau, 2019), and the mechanism supporting the ability to combine multiple tactile inputs has remained poorly understood.

A key neurophysiological mechanism supporting the ability to detect local differences in sensory input is lateral inhibition. Lateral inhibition implements a specific form of divisive normalization computation (Brouwer *et al.*, 2015; Rahman and Yau, 2019), in which the response of each unit is scaled by the response of a larger neural population, potentially enhancing contrast and local difference detection. Lateral inhibition can be approximated non-invasively by measuring evoked responses to digital stimulation and calculating response underadditivity (Gandevia *et al*., 1983; Hsieh *et al*., 1995; Ishibashi *et al*., 2000; Severens *et al*., 2010; Cardini *et al*., 2011, Cardini and Longo, 2016). Consistent with previous studies, we found increased underadditivity when participants were preparing to compare simultaneously delivered probe directions. In contrast, when participants were preparing to combine the two stimuli, underaddivity was significantly reduced. We argue that the strength of suppressive interaction between adjacent finger representations, which likely indexes somatosensory lateral inhibitory mechanism, is modulated top-down according to the perceptual task.

In a recent study, Canales-Johnson and colleagues (2020) found that whether participants perceived bistable auditory streams as one integrated stream or two distinct streams was reflected in the coherence of the neural activity within frontoparietal cortices. These results showed that integration vs. differentiation might be a global mode of coordination in fronto-parietal networks. Our study suggests that these putative modes are associated with different states of early cortical circuitry. Canales-Johnson *et al*.’s study relied on uncontrolled endogenous fluctuations in a bistable percept to switch between integrative/combining and distinct/comparison modes. Our study instead relies on strategic shifting, according to the current perceptual task. We speculate that higher cortical areas, such as frontoparietal networks, may be the source of the strategic signal that modulates early somatosensory cortical processing, adjusting the degree of inhibition, and thus the extent of observed underadditivity.

Our concept of distinct perceptual modes for integrative vs. discriminative processing recalls similar distinctions in the visual attention literature. For instance, Baek and Chong (2020) recently proposed two modes of processing in visual perception: ensemble perception, whereby observers extract a combined quality across multiple stimuli, and selectivity, whereby observers discriminate a specific stimulus among others. They explained the difference between these perceptual modes using a mechanistic model of selective attention. Distributed attention allows the brain to extract the mean activity across a population of sensory neurons, whereas focussed attention narrows the activity profile down to a smaller population. Focussed attention might achieve this selection of a smaller subset of sensory neurons by increasing lateral inhibition to provide tighter tuning.

A second mechanism that may regulate the balance between integration and discrimination is divisive normalization. This has been considered a canonical neural computation (Carandini and Heeger, 2012). During divisive normalization, the response of a single unit is divided by the response of a population. This has a similar net effect to lateral inhibition, since it again emphasises local departures from the population mean, but it does not involve the explicit mechanism of inhibitory interneurons associated with lateral inhibition in the visual system. Likewise, several studies suggest that engaging attentional mechanisms the brain can control the parameters of divisive normalization (Reynolds and Heeger, 2009; Brouwer *et al*., 2015).

However, our tasks always required processing information from both digits. In comparison task, participants had to report the exact difference between the stimuli, whereas in comparison task they had to report the exact average between the stimuli. The pattern of results in the combination task confirmed that participants did successfully divide their attention between digits, rather than merely attending selectively to one digit (see Figure 2B). What differed between the tasks was the way in which information from one digit was related to information from another. Our results suggest that the neural circuitry of sensory systems can potentially be tuned to implement either of two perceptual modes (extracting differences or extracting overall features), without engaging distinct attentional mechanisms. Selecting which operation is performed may involve adjusting inhibitory links, normalization pools, or both.

The focus of the present study was the P40 component, because it is considered a marker of S1 processing (Allison *et al.,* 1989). In addition, inter-finger suppression has not been found to affect earlier components such as N20 (Forss *et al.,* 1995). However, a recent study showed that important trial-by-trial variability dynamics occur as early as 20 ms after tactile stimulus onset (Stephani *et al.,* 2020). Our 30 Hz low-pass filter did not allow us to assess very early components. Therefore, we additionally re-processed our data without any low-pass filter, to maximise the opportunity of detecting early components (please see S1 of supplemental material). Indeed, a clear N23 component was identifiable in the unfiltered data, approximately covering a 20 – 25 ms time-window. However, a 2−2 ANOVA analysis analogous to the one performed on the P40 component did not reveal a significant interaction between task and number of fingers stimulated (*F*_1, 14_ = .55, *p* = .47, *η*_*p*_^*2*^ = .04), suggesting that earlier components such as the N23 do not display the task-specific modulations of somatosensory suppression that we specifically found for the P40. In contrast, this task-specific modulation of suppression remained significant for the P40 component when no low-pass filter was applied (*p* = .02).

One previous study reported multisensory modulation of somatosensory suppressive interactions within P40 time-window by simply viewing one’s own body (Cardini *et al*., 2011). That finding already suggested that the strength of subadditivity that indexes lateral inhibition may not be constant, but can be modified by other factors. However, to our knowledge, the wider question of how and why lateral interactions might be adjusted has rarely been considered. Studies of olfactory processing in animals assume that such interactions always aim at maximum acuity (Yokoi *et al*., 1995), providing enhanced pattern separation for specific molecules within complex mixtures. However, one recent study suggests that the circuitry underlying pattern separation is plastic, and shaped by experience of perceptual discrimination (Chu *et al*., 2016). A study of drosophila visual system (Keleş and Frye, 2017) found that blocking GABAergic inhibition resulted in reduced visual responses to a single moving object, and increased responses to wide-field pattern motion. Yet, the results of this study can be interpreted in relation to attentional focus either to local variations or to overall gist.

Our study goes further, in suggesting that the degree of lateral interaction can be strategically engaged, as a distinct mode of perceptual processing, according to the requirements of a task. When participants need to favour differentiation based on specific details, increased inhibition may amplify small local differences. In contrast, when participants are preparing to access an overall synthesis of complex inputs, reduced inhibition may facilitate aggregation and generalisation. Our results emphasise the potential flexibility of tuning neural circuitry, a process that may be crucial for successful interaction with the world. For example, Bertone and colleagues (2005) speculated that the increased ability to detect and distinguish individual stimuli and reduced integrative processing seen in individuals with autism might be due to unusually strong lateral inhibition (also see Gustafsson, 1997). If this is so, our results raise the intriguing possibility that inhibition might not be excessive *per se*. Rather, autistic individuals might show reduced flexibility in adjusting inhibitory local networks when the task requires it. More generally, Herz and colleagues (2020) recently suggested that healthy thinking and perception is characterised specifically by the degree of flexibility to continuously change across a continuum spanning from ‘narrow’ states of mind based on bottom-up processing, to ‘broad’ states based on top-down processing. Our results also suggest that the cognitive flexibility of tuning neural circuitry responsible for sensory cortical interactions may play a key role in shaping how we experience the world around us.

## Supporting information

Main measures

## Acknowledgements

This study was supported by a research contract between NTT and UCL, and an MRC-CASE studentship MR/P015778/1.

## Supplemental material

### S1. Analysis of somatosensory evoked potentials without low-pass filter to identify early components

The main manusript shows data that was low-pass filtered at 30 Hz, which could have concealed the rapid early components discussed in the MNS literature (e.g., Desmedt et al., 1983; Sehm et al., 2013). We therefore re-ran our EEG preprocessing without any low-pass filter to identify whether there were any earlier components. We did then find a clear early component in the 20 – 25 ms time-window with a peak at 23ms (−.35 mV at the peak). We ran a similar analysis on this N23 component as for the P40 component (see main manuscript) to investigate whether there was task-related modulation of suppression.

As for the P40, we used a repeated-measures ANOVA with factors task (combination vs. comparison) and stimulation (both vs. index + middle). The analysis did not yield a significant interaction (*F*_1, 14_ = .55, *p* = .47, *η*_*p*_^*2*^ = .04), suggesting no modulation between the tasks for the N23. For consistency, we also analysed the unfiltered P40 component, but starting from 25 ms rather than 20 ms to not overlap with the N23 component. The analysis yielded a significant interaction (*F*_1, 14_ = 6.7, *p* = .02, *η*_*p*_^*2*^ = .32) with simple effects showing a significant suppression in comparison (*p* = .002), but not in combination task (*p* = .72).

---

**Figure S1.**
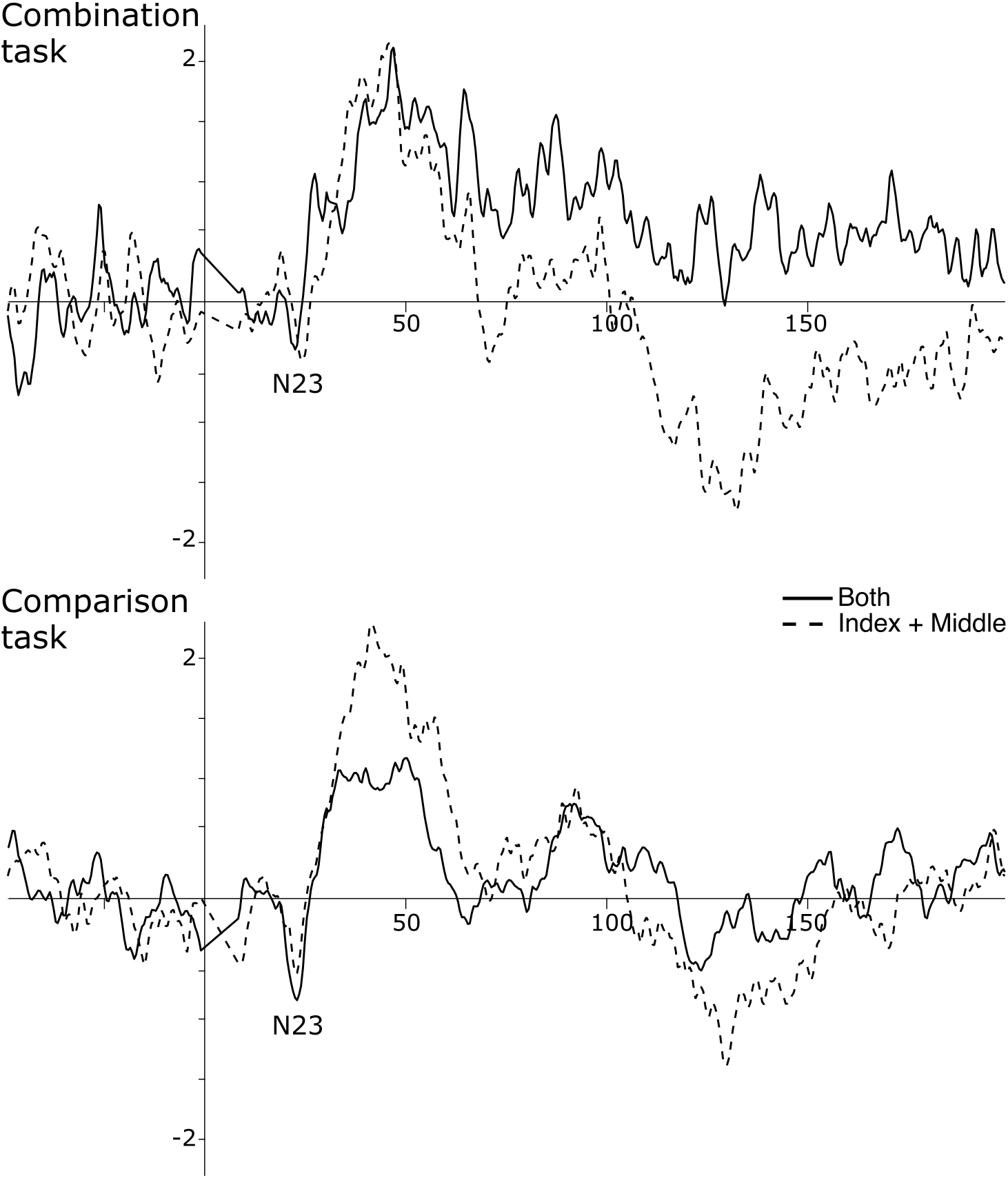
Somatosensory evoked potentials without low-pass filter. Upper panel displays combination task while lower panel displays comparison task. Solid line represents grand-average of trials were both fingers (index & middle) were simulated simultanously. Dashed line shows grand average of sum of individual simulations, which represent the predicted amplitude under the assumption of no supression. The waveforms represent pooled activity across contralateral somatosensory electrodes (C5, C3, CP5, CP3).

